# High-throughput screening of high protein producer budding yeast using gel microdrop technology

**DOI:** 10.1101/830596

**Authors:** Hirotsugu Fujitani, Soichiro Tsuda, Tomoko Ishii, Masayuki Machida

**Affiliations:** Biomedical Research Institute, National Institute of Advanced Industrial Science and Technology (AIST), Ibaraki, Japan; On-chip Biotechnologies, Tokyo, Japan; Genome Biotechnology Laboratory, Kanazawa Institute of Technology, Ishikawa, Japan

**Author notes:** Corresponding author: Masayuki Machida.

## Abstract

The need for protein production has been growing over the years in various industries. We here present a high-throughput screening strategy to isolate high producer budding yeast clones from a mutagenized cell population using gel microdrop (GMD) technology. We use a microfluidic water-in-oil (W/O) emulsion method to produce monodisperse GMDs and a microfluidic cell sorter for damage-free sorting of GMDs by fluorescently quantifying secreted proteins. As a result, this high-throughput GMD screening method effectively selects high producer clones and improves protein production up to five-fold. We speculate that this screening strategy can be applied, in principle, to select any types of high producer cells (bacterial, fungal, mammalian, etc.) which produce arbitrary target protein as it does not depend on enzymes to be produced.

## Introduction

The need for protein production has been growing in recent years, owing to the rapid advancement of biopharmaceuticals such as therapeutic antibodies (1). Industrial enzymes have been a major use of protein production, widely used in various industries, such as food, fuel, and pharmaceutical industries. As this need is expected to increase further (2), more efficient protein production is required to cope with the growing need.

Microbes have been serving as a workhorse for protein production for a long time because of ease of genetic engineering and the fast growth. It is, however, well recognized that a microbial population in general shows metabolic heterogeneity, in which individual cells show different protein expression levels due to transcriptional noise (3). Thus, selection of high producing subpopulation is a crucial problem when producing proteins more efficiently at the industrial scale (4).

A number of selection methods have been developed so far. The most widely used (thus conventional) technique is the limiting dilution method, where cell population is diluted in well plates until single cells are isolated in individual wells, followed by subsequent protein quantification assays, such as enzyme-linked immunosorbent assay (ELISA). This traditional method is labor-intensive, time-consuming and low-throughput, thus alternative high-throughput screening (HTS) methods have been actively sought after.

Florescent-activated cell sorter (FACS) is an alternative HTS method for isolation of high-producing cells (5,6). For example, high producer cells were isolated by FACS based on fluorescent intensity of green fluorescent protein (GFP), co-transfected with a target protein (7). However, there is a general trade-off between the protein productivity and growth rate due to metabolic burden imposed by heterologous protein production.

One of the alternative HTS methods that circumvents the trade-off issue is a gel microdrop (GMD) method (9). Individual cells are encapsulated into agar GMDs and cultured to form colonies and secrete target proteins within. The proteins are confined in the GMDs due to limited diffusion of molecules or by cross-linking to gel materials (e.g., by avidin-biotin interaction) (6). Captured proteins are fluorescently labelled in order to link protein production and fluorescence intensity. This method prevents users from selecting high-producing but slow-growing cells because the production level is assessed by the total amount of target protein secreted by a group of producer cells.

In this paper, we set out to address two issues pertaining to GMD-based screening method. First, the conventional method for producing GMDs create polydisperse GMDs ranging from tens of microns to sub-millimeter in diameter. Larger GMDs need to be filtered out to avoid GMDs clogging inside FACS. This means some portion of whole yeast population contained in the large GMDs will be lost at this step, which effectively decrease the size of entire yeast population to be screened. Plus, the method requires a large volume to produce GMDs at a time (typically 10 mL), which is costly and hence makes it difficult to test various experimental conditions. We overcome these issues by creating monodisperse GMDs using microfluidic droplet generator (Supplementary Fig. 1). This method typically requires tens of hundred microliters and uniform-size GMDs eliminates the need of filtering prior to sorting.

Second, cells sorted by cell sorters can die or show little growth after sorting because of sorting-induced cellular stress (8,10,11), which is also the case with GMD-based cell sorting. To improve the viability of sorted cells, we employed a microfluidics-based cell sorter, which cause much less damage or stress to the cell, and hence show better viability.

By combining these two features, we show that GMD-based yeast screening improves the protein yield up to five-fold compared to the original strain only in one round of screening.

## Materials and Methods

### Construction and cultivation of luciferase-producing budding yeast BY4741 strain

A plasmid used in this research (Figure 1B) was prepared by combining the vector DNA and the fragments amplified by PCR using Gibson Assembly (New England BioLabs). The vector harbored the *URA3* and the *leu2-d* markers and the 2-m replication origin derived from the pYEX-S1 (Clontech) backbone. The protein expression cassette consisted of the *GAL1* promoter, secretory luciferase and the *CYC1* terminator. The prepro-alpha-factor leader peptide of *S. cerevisiae* was fused to *Metridia longa* luciferase derived from pMetLucReporter (Clontech) after removal of its original signal peptide and was further fused to Halo-tag derived from HaloTag Control Vector (Promega) at the C-terminus. The FLAG and the Hisx6-HA tags were introduced directly at the downstream of the prepro-alpha-factor leader peptide and the luciferase, respectively. Transformation of yeast BY4741 strain was conducted according to a standard protocol of *S. cerevisiae* Direct Transformation Kit *Wako* (Fujifilm Wako Chemical, Osaka, Japan).

**Figure 1.**
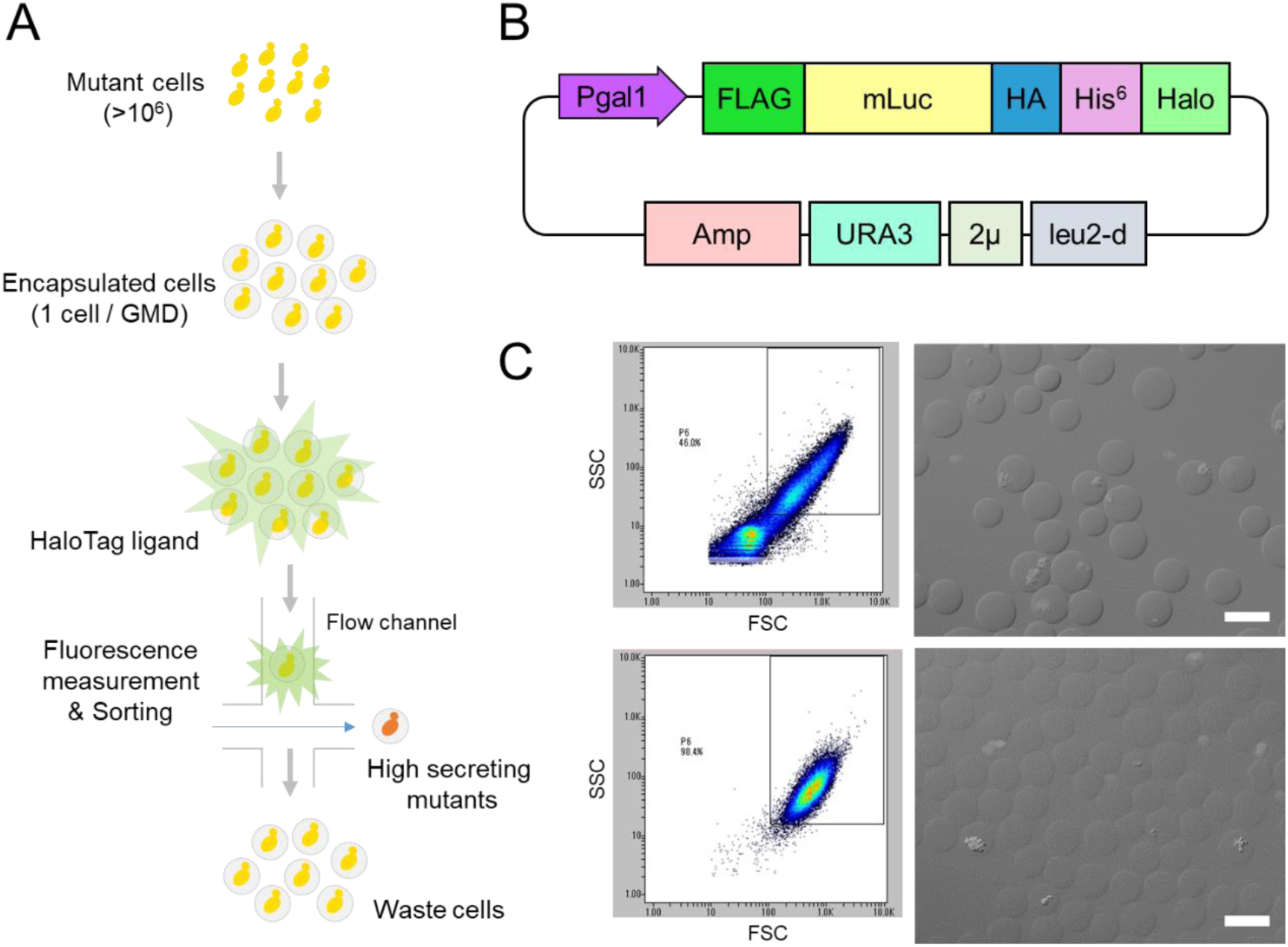
(A) Schematic illustration of the high-throughput screening workflow for high protein producing yeast cells. (B) Plasmid map of a plasmid producing mLuc with gal1 promoter used in this study. (C) Comparison of GMD formation method: Conventional membrane filtration method (upper row) and microfluidic droplet method (lower row). Left column shows dot plots of forward scatter (FSC) and side scatter (SSC) of GMDs analyzed by the microfluidic cell sorter. Right column shows microscope images of GMDs.

The transformant was grown in the medium containing 0.67% Yeast Nitrogen Base w/o Amino Acid (DIFCO) supplemented with –Ura DO Supplement, 100 mM sodium phosphate (pH 7.0) and 2% carbon source (glucose or galactose as indicated in the text).

### UV mutagenesis of budding yeast BY4741 strain

The transformed BY4741 stain was exposed to UV light to introduce random mutagenesis in the genome to screen high producer mutants using cell sorter. To do this, the yeast cells grown on SD-ura medium containing glucose were first diluted to 1.0 × 10^6^ cells mL^-1^ with SD-ura medium containing galactose, then pipetted on a sterile plastic surface (10 µL×30 spots). They were irradiated by UV light for 0 to 120 seconds. After UV exposure, the yeast suspensions were collected in a tube for GMD encapsulation.

### Microfluidic generation and cultivation of GMDs

Mutagenized yeast suspension was mixed with 2.5% molten low-melting point agarose gel at 4:1 volume ratio. The suspension-agar mixture was loaded into a sample well of a DG800 cartridge (On-chip Biotechnologies, Tokyo, Japan) to generate water-in-oil (W/O) emulsion using On-chip droplet generator (On-chip Biotechnologies, Tokyo, Japan). Two percent 008-FluoroSurfactant in HFE 7500 (RAN Biotechnologies, USA) was used as the continuous oil phase. The pressures of cell suspension and oil were maintained at 30 kPa and 20 kPa, respectively, to keep the size of generate W/O emulsion around 50-60 µm. The whole droplet generator unit was kept in a temperature control unit at 37°C to prevent the agarose from solidifying and form stable-size droplets. Three hundred microliter of the yeast suspension was encapsulated into the emulsion for each sample. The W/O emulsion was kept on ice for at least 30 min to make GMDs by solidifying the agarose gel. The oil phase with the fluorinated surfactant was removed by adding 10% 1H,1H,2H,2H-Perfluoro-1-octanol (Sigma-Aldrich) in HFE 7500 (14) and GMDs were suspended in SD-ura medium containing galactose.

GMDs containing yeast cells were cultivated with a shaking incubator at 30°C and overnight.

### Fluorescence staining of GMDs

To stain proteins secreted from cells within GMDs, Halo Tag Alexa Fluor 488 Ligand (Promega) was diluted with moderate amount of PBS. The diluted Halo tag ligand were mixed with cultivated GMDs at room temperature and further incubated for 30 min. After washing three times with PBS, the stained GMDs were observed with fluorescence microscope.

### Sorting and microscopic observation of GMDs

Sorting of GMDs containing high protein producer yeasts was performed using On-chip Sort (On-chip Biotechnologies, Tokyo, Japan). On-chip Sort employs a microfluidics-based sorting mechanism with disposable microfluidic chip (Supplementary Figure 2). Eighty micron channel width disposable sorting chip (Z101, On-chip Biotechnologies) was used for cell sorting with On-chip T buffer as sheath liquid. The sample was flown through a microfluidic channel at approximately 300-500 events per second and in total 200-500 target GMDs were sorted. After sorting, collected GMDs containing yeast were observed using differential interference contrast and fluorescent microscope (BX3-URA, Olympus, Tokyo, Japan) for morphological analysis of yeasts.

### Luciferase assay of sorted cells

Sorted GMDs were streaked onto agar plates containing the SD-ura medium with galactose for further cultivation and colony formation. Each colony was picked and suspended into 2 ml of SD-ura medium containing galactose at pH7.0. The suspensions were incubated with shaking at 30°C, 150 rpm for 24 hours. The supernatant was retrieved and applied to luciferase assay. The luciferase assay was conducted according to a standard protocol of Ready-To-Grow Dual Secreted Reporter Assay (Clontech Laboratories, Inc., US) except that the amount of substrate was reduced to half of the defined amount.

## Results

### Workflow of high-throughput GMD screening for high producer mutant cells

First, we describe a workflow of our screening method for high protein producer cells using GMD and cell sorter (Fig. 1A): A plasmid with mLuc gene and *gal1* promoter (Fig. 1B) were transformed into yeast cells and mutagenized by UV exposure. The mutant yeast cells were diluted to ∼1×10^6^ cells/mL and encapsulated in agarose gel using microfluidic droplet generator so that most likely only one cell would be embedded in one GMD (i.e., Poisson parameter λ=0.1). GMDs including mutant cells were incubated overnight and then luciferase secreted in the GMDs were stained by HaloTag Alexa Fluor 488 ligand. GMDs with strong fluorescence were sorted by a microfluidics-based cell sorter because strong fluorescence indicates more protein production and secretion. Sorted GMDs were cultured on agar plates to form colonies. Each colony was picked up and sub-cultured with nutrient medium for luciferase assay.

### Comparison of GMD size formed by different GMD formation methods

Prior to sorting of GMDs, we investigated the effect of different formation methods on the size of GMDs. Figure 1C shows dot plots and microscope images of GMDs containing yeast cells grown overnight. The dot plots show forward scatter (FSC) and side scatter (SSC) obtained by the microfluidic cell sorter, which represents the size and the internal complexity of samples, respectively. GMDs formed by a conventional membrane filtration method (16) show a wide distribution of points in the dot plot (Fig. 1C upper left) whereas those by the microfluidic droplet generator did much narrower distribution (Fig. 1C lower left). This indicates the latter samples are uniform in terms of size and internal structure, compared to the former ones. Indeed, microscopic images confirm this observation: The size of GMDs made by the microfluidic method was monodisperse, while the one by the conventional method varied even after filtration by a 70 µm cell strainer. Furthermore, the microfluidic method does not require filtration and thus the whole GMDs generated can be used for screening.

### Sorting of GMD and Sorting

The mutagenized yeast population encapsulated in GMDs was grown overnight at 30°C 150 rpm, then applied to microfluidic cell sorter, On-chip Sort. The amount of produced proteins was quantified by fluorescent ligand (HaloTag Alexa Fluor 488 ligand) covalently bound to HaloTag conjugated to mLuc. The fluorescent ligand is expected to label proteins secreted out of cells because it is a cell membrane impermeable compound. We primarily focused on FL2 (detection wavelength: around 575 nm) and FL3 (detection wavelength: around 620 nm) fluorescence channels on On-chip Sort because of the fluorescent ligand. The dot plot of FL2 against FL3 fluorescence typically showed a distribution with two long tails expanding towards upper right (Fig. 2A). From microscope image analyses of sorted samples, we found that the upper tail consisted of small contaminants (e.g. small fibers or plastic pieces with autofluorescence). In contrast, the lower tail consisted of GMDs containing budding yeast cells. We found that small colonies were typically formed within GMDs (Fig. 2B and C). We split the long tail into three segments, named as P7, P8, and P9, based on the fluorescence intensity of FL2 channel. In P7, we found that some of the sorted samples showed strong fluorescence despite the colony size (Fig. 2B red circle). Considering that they were small in size or did not form any colonies, we speculated that they were dead cells. They can be false positive samples because a mass of mLuc proteins released out of the loose cell wall were stained by fluorescent HaloTag ligand. On the other hand, GMDs sorted from the P8 segment were observed to show moderate fluorescence with growing colonies found within GMDs (Fig. 2C). GMDs from P9 segment also contained similar colonies, but with less fluorescence. For these reasons, we decided to sort samples from P8 segment. Typically, around 1000 samples in one experiment were sorted with P8 gate and cultured for further analysis.

**Figure 2.**
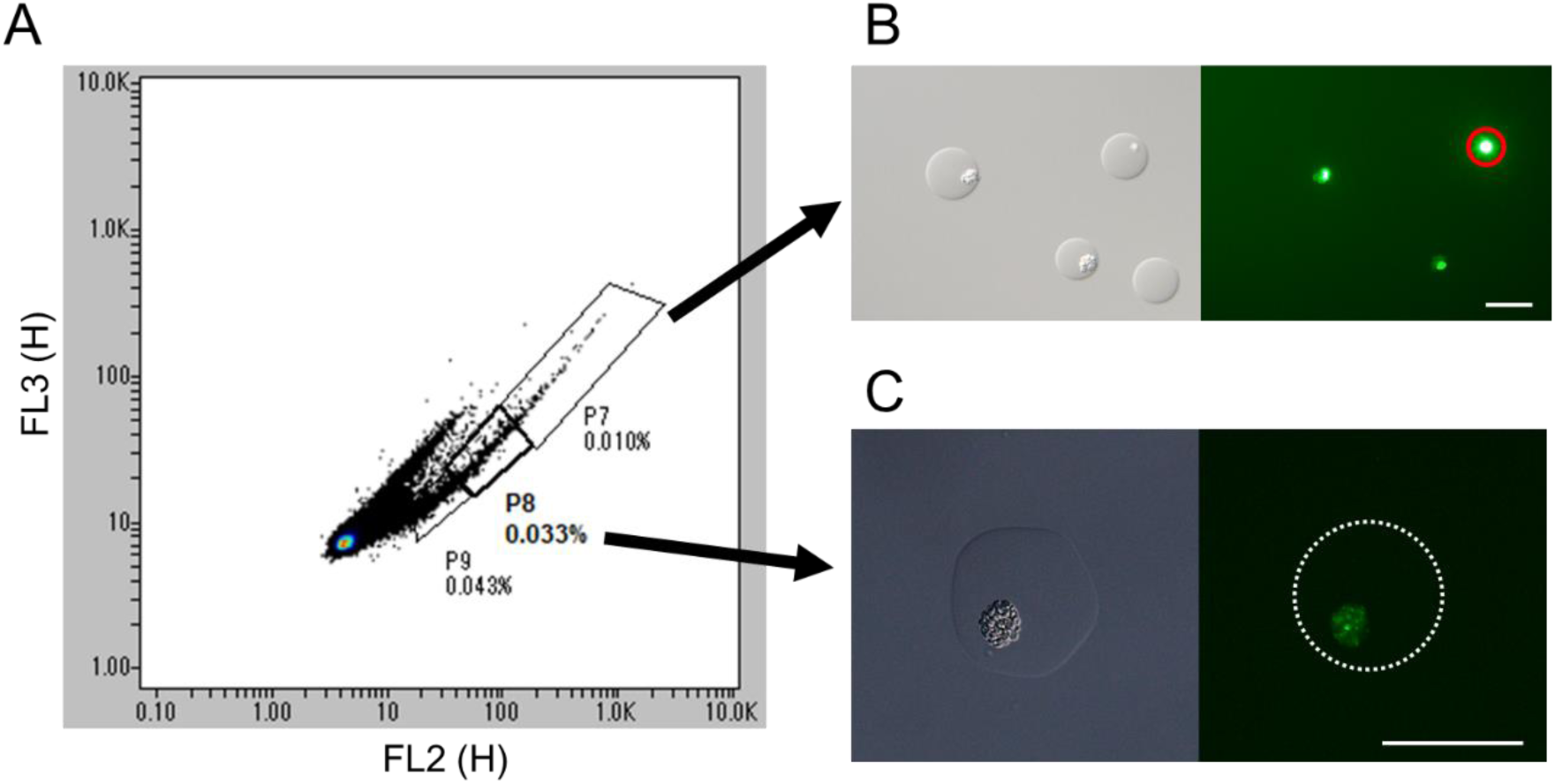
(A) An example dot plot of FL2 against FL3 (peak height, H) for sorting of GMDs. (B) Microscope images of sorted GMDs from P7 segment: Differential interference contrast (left) and green fluorescence (right). Red circle indicates dead cell(s). (C) Microscope images of sorted GMDs from P8 segment. Dotted line shows border of GMD.

### Luciferase assay

GMDs sorted from P8 segment were sub-cultured on agar plates containing the SD-ura medium at 30°C for at least four days until colonies were visible. Colonies on the plates were individually transferred 96 well plates with liquid SD-ura medium with galactose and cultured overnight. The supernatant of total 14 sorted samples as well as the original strain was applied to luciferase assay. A half of all sorted samples indicated higher protein producing activity than the original strain, of which one sample (sample P8-7) showed more than twice activity and another sample (sample P8-12) was five-fold higher (Fig. 3).

**Figure 3.**
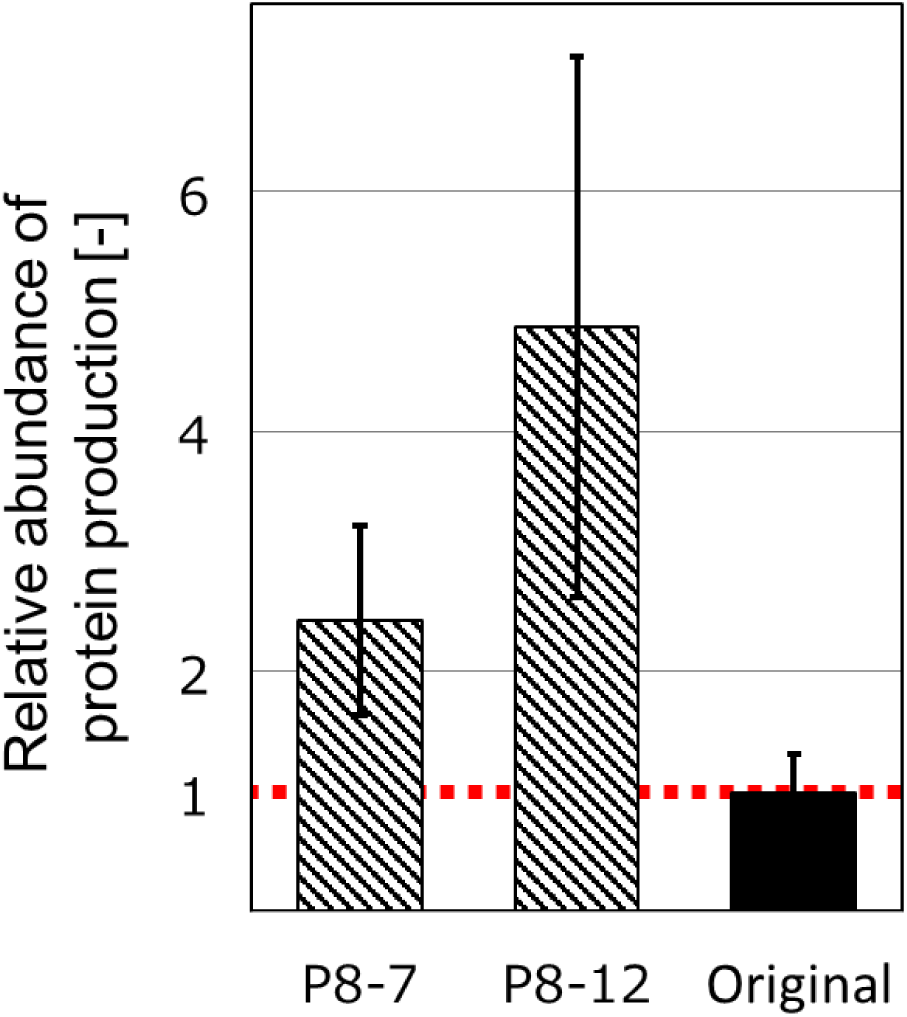
Relative abundance of protein production based on the luciferase assay of high producing cells. Cells sorted from P8 area were compared with original cells. Error bars show standard deviation (*n* = 3, technical triplicates).

## Discussion and conclusion

We have shown that, as a proof-of-concept, our GMD method effectively selects high producer clones and improves protein production up to five-fold from only one round of selection. This work combines microfluidic GMD generation and flow cytometry for HTS. Similar work using microfluidics and GMD has been done in recent years, such as selection of oil-producing microalgae (12) and directed evolution of xylanase-producing yeast (13). As our selection strategy does not depend on enzymes to be produced, in principle it can be applied to select any types of high producer cells (bacterial, fungal, mammalian, etc.) which produce arbitrary target protein. We also speculate that the strategy can be applied to the selection of high producer non-model organisms for which genetic engineering cannot be used. This can be possible, for example, by labelling target proteins by fluorophore-conjugated antibody or by linking the activity of secreted enzymes with signal intensity using fluorescent probes based on Föster resonance energy transfer (FRET). We foresee a wide range of applications for selecting high producer cells, as this method is capable of sorting not just microbes, but also mammalian cells which are relatively prone to damage or stress by cell sorting.

## Acknowledgement

This work was supported by Kanazawa Institute of Technology (M.M.) and Leading Initiative for Excellent Young Researchers (LEADER) program (S.T.).

